# Global analysis of multi-mutants to improve protein function

**DOI:** 10.1101/2020.12.03.408732

**Authors:** Kristoffer E. Johansson, Kresten Lindorff-Larsen, Jakob R. Winther

## Abstract

The identification of amino acid substitutions that both enhance the stability and function of a protein is a key challenge in protein engineering. Technological advances have enabled assaying thousands of protein variants in a single high-throughput experiment, and more recent studies use such data in protein engineering. We present a Global Multi-Mutant Analysis (GMMA) that exploits the presence of multiply-substituted variants to identify individual amino acid substitutions that are beneficial for the stability and function across a large library of protein variants. We have applied GMMA to >54,000 variants of green fluorescent protein (GFP), each with known fluorescence output, and each carrying 1–15 amino acid substitutions. The GMMA method achieves a good fit to the data while being analytically transparent. The six top-ranking substitutions are demonstrated to progressively enhance GFP and in general, our analysis recovers nearly all the substitutions previously reported to be beneficial for GFP folding and function, only using a single experiment as input.

**Significance Statement:** Protein engineering is carried out to improve proteins for practical applications by changing one or more amino acid residues in a protein. We present a method termed global multi-mutant analysis (GMMA) that helps solve two problems in protein engineering. First, because many proteins are already highly optimized, it can be difficult to identify individual variants that further improve function. Second, while it is possible to combine variants with small individual effects, such approaches may be hampered by non-additivity. GMMA identifies combinable effects of single substitutions from a large set of variants each carrying multiple substitutions. We demonstrate the approach on a set of 54,000 variants of green fluorescent protein and identify many enhancing single-substitutions from a single experiment.

## Introduction

A major challenge in practical applications of proteins is the engineering of protein stability while at the same time maintaining the function of the protein. New developments in biotechnology are continuously applied to address this challenge with high-throughput methods currently in focus. Synthesis, screening and sequencing may today be performed for thousands of protein variants in parallel via *Multiplexed Assay of Variant Effects* (MAVE) also known as *deep mutational scanning* [1, 2]. Such experiments can identify loss-of-function variants with high accuracy, but are often unable to gauge more subtle effects. For example, stabilizing or mildly destabilizing substitutions are likely to have a minor, if any, detectable impact on protein function and are therefore more difficult to identify from such experiments. Additive contributions to stability or catalysis are particularly attractive for protein engineering [3,4] and the ability to extract such from MAVE experiments could impact on the field substantially. Stabilizing and folding-enhancing substitutions may also provide robustness to other mildly destabilizing substitutions and thus enhance evolvability towards novel functions [5].

When a protein library contains a high fraction of variants with two or multiple amino acid substitutions, statistical models have been used to investigate how the effect of individual substitutions combine. One important technical consideration is the mathematical framework to capture (non-)additivity of the variant effects [6]. For example, one would often find that the thermodynamic stability effects of two substitutions are additive when quantified by free energy differences, whereas the corresponding equilibrium constants would not be additive, but rather multiplicative. Thus, more general models typically consider additive single-substitution effects, here generally referred to as a protein fitness potential, and transforms the sum of these to the assayed quantity via various non-linear functions to describe the data generated by a MAVE [7–9]. One study further showed that a thermodynamic model could be used to quantify the effects of single-substitutions on protein binding and structural stability from experimental measurements of the effects of both single and double mutants on binding [9]. This model was later shown to capture structural stability accurately [10] and a similar approach was successfully applied to fit deleterious effects of multiply-substituted variants [8]. The fitness potential has some analogy to free energies or statistical potentials [11].

Here, we present a method, Global Multi-Mutant Analysis (GMMA), to estimate fitness potentials, that enable the identification of amino acid substitutions that have a general enhancing effect with little dependency on the sequence context, and thus substitutions with the potential to be additively combined. We demonstrate that these single-substitution effects may be informed by variants with multiple amino acid substitutions, here referred to as multi-mutants. The central idea is to identify beneficial substitutions by their ability to compensate other substitutions that decrease function for example by destabilizing the protein. For example, while a stabilizing substitution may not have a measurable effect in the background of an already stable protein, it can be identified in the background of one or more destabilizing substitutions that together cause decreased activity of the protein. The concept is similar to the “partner potentiation” [12] or “disrupt-and-restore” [13] principles formulated previously, but here generalized to large libraries of diverse multi-mutants.

We have applied our analysis to an experiment that reports the fluorescence of >54,000 variants of green fluorescent protein (GFP) each containing 1-15 of the total of 1,879 unique single-amino-acid substitutions observed in the experiment [7]. The GFP variants were generated using error-prone PCR (epPCR) and expressed in fusion with a red-fluorescent protein to correct for variations in expression levels. Then, fluorescence activated cell sorting (FACS) was used to divide the cells into eight fractions based on their level of green fluorescence, and each fraction was sequenced to link an intensity of fluorescence with a genotype. We have applied GMMA to the results of this experiment to estimate the effects of 1,107 single-substitutions, and identify substitutions that are beneficial for both function and stability and thus directly applicable to engineering studies. As validations of the approach, we show that the model accurately predicts experimental variant effects that were not used in training, and show that the top six substitutions may progressively be combined in new GFP variants that achieve a more than 6-fold enhancement of cellular fluorescence compared to the starting point when expressed in *E. coli* cells. Finally, we show that the method identifies a large number of substitutions that have previously been described to be beneficial for improving GFP.

## Results and Discussion

### Concept

Before formulating the fitness potential, we briefly describe the qualitative concept that inspired the development of GMMA (Fig. 1). While exploring the GFP data for information on beneficial mutations, we considered the ability of the functional protein to tolerate random mutations before losing function (Fig. 1e). This showed that the “wild type” (or reference sequence) can tolerate approximately four average amino acid substitutions before the average brightness drops to 50%. With this as a reference, we next considered the subset of variants that carries a specific substitution, e.g. V163A, and find that in this background the protein can tolerate on average an additional five substitutions before the average brightness drops to 50% (Fig. 1f). Thus, we concluded from this simple analysis that V163A possesses some enhancing property across all sequence contexts in the subset of GFP variants.

**Figure 1.**
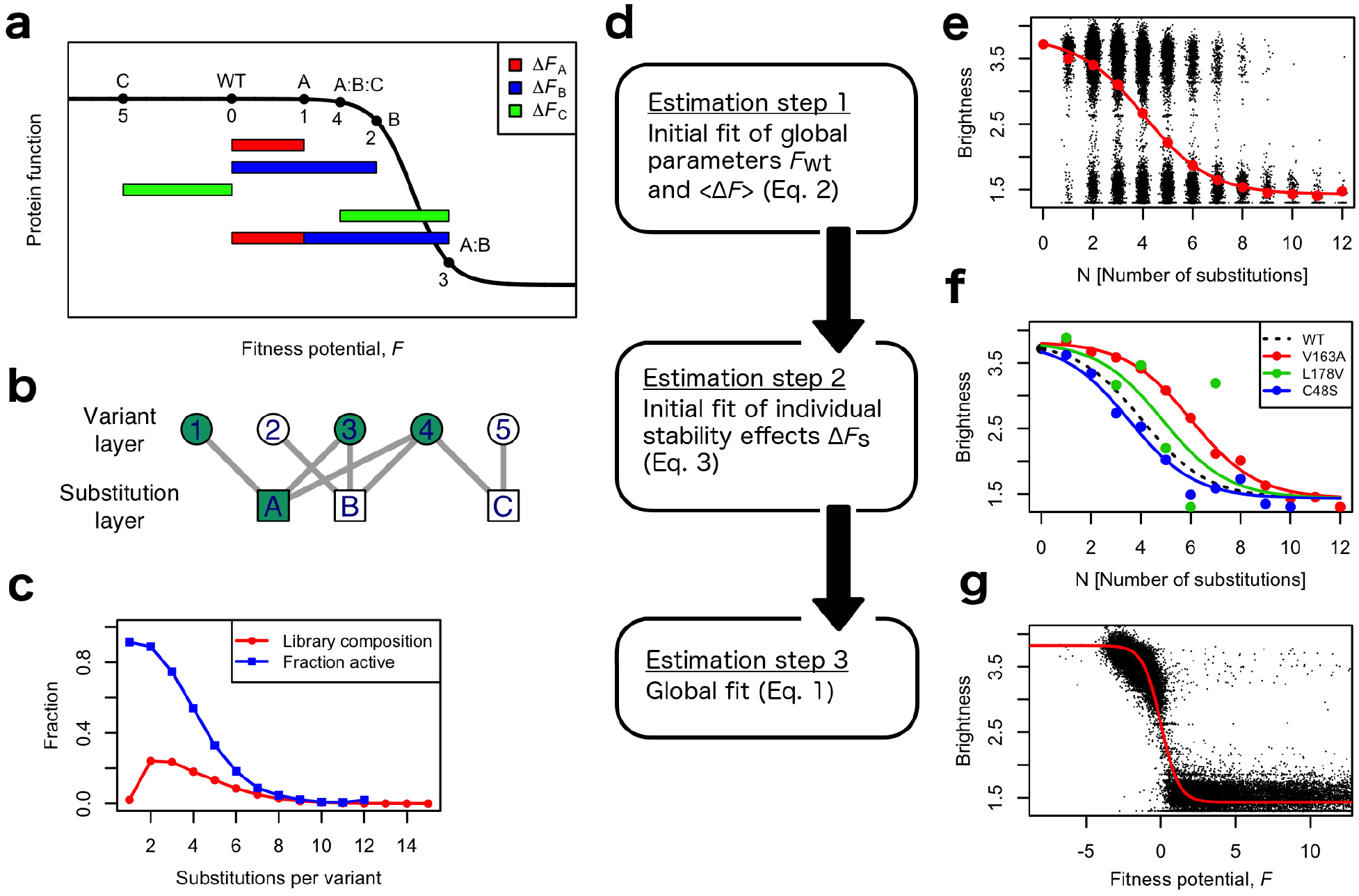
**(a)** A schematic outline of the GMMA approach. Consider a protein (WT) with five variants (named 1–5) that are composed of one or more of three substitutions (named A–C). The lengths of the colored bars represent the magnitude of the additive effects of these three substitutions. All single-substitution variants show wild-type-like activity, with the most functionally abated variant, B, only being slightly less active than the wild type. While both variants A and B are active, the double mutant A:B is inactive. Thus, we infer that, in an additive model, A and B both decrease the fitness potential. Substitution C on its own does not appear to affect activity. However, when it is introduced into the inactive A:B background (to form A:B:C) it is able to compensate the loss of function, and we thus infer that substitution C enhances the fitness potential with the magnitude of the green bar. (**b**) The five variants and the three substitutions may be represented in a bipartite graph, and as an example we highlight (in green) the subset of protein variants used to estimate the effect of substitution A. (**c**) Multi-mutant composition of the GFP library (red) shown together with the fraction of active variants (blue). The high population of variants in the transition region between 2 and 6 amino acid substitutions makes this excellent for GMMA. **(d)** Three-step estimation strategy. The three steps relate to panels e, f and g respectively **(e)** Fit of initial global parameters (red line) to the average brightness of variants with a given number of substitutions (red points) using Eq. 2. The GFP library is shown with random jitter in the horizontal coordinate to ease visualization (black dots). **(f)** Three examples of initial fits of individual stability effects (colored lines) compared to the WT fit from panel e (dashed line). Each Δ*F_s_* is fit to the average *N*-mutant brightness (points) calculated only using the subset of variant that contains *s*. Substitutions that are mildly detrimental to the fitness potential, like C48S, lowers the endurance towards additional mildly detrimental substitutions and thus the system inactivates with fewer substitutions and the curve shifts left (blue line) compared to the wild-type (dashed line). On the other hand, enhancing substitutions, like V163A, may be identified by their ability to increase the endurance towards substitutions, thus shifting the curve to the right (red line). Substitutions observed in few variants are difficult to fit, here demonstrated with L178V observed in only 14 variants compared to 677 and 194 variants for V163A and C48S respectively. **(g)** Result of the final global optimization (red line) to all variants (black dots).

We formulate this concept by an analogy with traditional measurements of protein stability [14], and describe the inactivation as a two-state process where the system, a “variable protein”, is in equilibrium between an active and inactive state. This inactivation may be probed by mutating the protein and substitutions with the ability to rescue activity of an inactive variant, are identified as stabilizing or generally enhancing (Fig. 1a). The equilibrium between the two states has an associated free energy, here referred to as the fitness potential *F,* which is linearly affected by the individual substitutions, *s*, that constitutes a variant, *v* ∈ {1… *V*}:

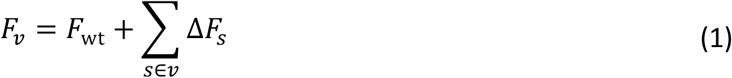

Here, *F*_wt_ is the fitness potential of the wild-type sequence and Δ*F_s_* is the effect of the substitution *s*. At fitness potential zero, the active and inactive states are equally populated and the assay readout is half of the maximal. A predicted assay readout is calculated from the fitness potential in analogy with the fraction of folded protein via Boltzmann probabilities (see Methods). Throughout this report, we will discuss the interpretation of the fitness potential in relation to conventional structural stability. We note that our approach is not limited to additive effects and that couplings may be included in Eq. 1, as long as the data warrants estimation of these. This formulation is similar to previous global models [6–9, 16,17], but here applied with the particular aim of identifying substitutions that enhance the protein in terms of the property that is assayed for.

The GMMA thus comprises a set of *V* equations, each describing the activity of a variant, and carrying a number of parameters, Δ*F_s_*, together with the global wild-type fitness potential, *F*_wt_, and baseline parameters. For such a system of equations, it is important to have more data (variants) than parameters (substitutions), which is possible with a multi-mutant library where a set of unique substitutions may be mixed in many different ways to make a larger set of variants. It is also important that all equations are coupled, and this may be tested by analyzing an undirected bipartite graph in which the protein variants constitute one layer of nodes (Fig. 1b, circles) and the unique substitutions the other layer (Fig. 1b, squares). This substitution-variant-graph formalism may be used to study many aspects of the multi-mutant library. For example, the degree distribution of variant nodes gives the distribution of the number of substitutions in the variants (Fig. 1c), which shows that most variants contain two to six substitutions. The fraction of active variants per *N*-mutant (Fig. 1c, blue line) shows that the substitutions in general decrease the fitness potential and that the fitness potential of wild-type GFP, *F*_wt_, is approximately 4 when measured in units of “general substitution effects”. GMMA identifies enhancing substitutions by compensation of negative effects, and it is important that the variant library has an *N*-mutant distribution that covers the transition region where the system loses activity.

The inactive state results from a number of deactivating amino acid substitutions, irrespective of mechanism, and is thus somewhat broadly defined. Some deactivating substitutions cannot be compensated by other enhancing substitutions, e.g. removal of a crucial functional side chain, and we term these *irreversible fatal substitutions* (IFS). IFS may be related to the assayed function of the protein, stability or folding hotspots, or for GFP, related to the chromophore maturation reaction. In principle, IFS have an infinite impact on the fitness potential but in practice they will be estimated to an arbitrary high value. This highlights an important difference from conventional structural stability and a distinct advantage that GMMA only identifies substitutions that support the assayed function.

In the interpretation of the GFP data it is important to realize that the deactivation-via-substitutions is qualitatively different from conventional *in vitro* unfolding experiments where unfolding is induced after the irreversible chromophore maturation. Indeed, one of the best-known enhanced variants of GFP is named *superfolder GFP* (sfGFP) and this name refers to the ability to fold and mature before creating non-fluorescent inclusion bodies [18]. For GMMA, the considered equilibrium between the active and inactive states, may likewise lie *before* the maturation reaction with only the active state being able to mature. Furthermore, the physical situation may closer resemble a steady state where active protein is removed from the equilibrium by irreversible maturation, possibly in a chaperone dependent fashion [19], and inactive protein is removed, e.g. by protease degradation or aggregation. In this scenario, maturation kinetics could also influence the outcome of GMMA.

### Model estimation

The global fit of the effects of the individual substitutions is complex for at least two reasons. First, the global parameters including baselines (Fig. 1a, black line) are estimated simultaneously with the fitness potential of all individual variants (Fig. 1a, abscissa values). The trade-off between adjusting the curve or the points may result in a rough optimization surface featuring many local optima. Second, many substitutions are observed in only few variants and may be poorly determined with greater uncertainties that are combined with otherwise well-determined substitution effects. To address these complexities, we use three-step estimation procedure that relies on initial estimates of individual effects (Fig. 1d). The first two steps focus on achieving initial estimates of global parameters (Fig. 1e) and individual effects (Fig. 1f) using a mean-field approach. The final global optimization in step three enhance consistency by allowing the fit to converge to the nearest optimum (Fig. 1g).

In the first step, an initial global wild-type potential, *F*_wt_, is estimated together with the average effect of a substitution, 〈Δ*F*〉 (Fig. 1e). This average effect replaces the sum in Eq. 1:

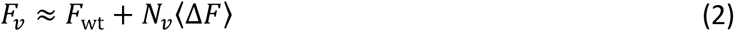

where, *N_v_* is the number of substitutions in variant *v*. While this assumption is rather crude, it is only used in the estimates of initial values in step two, and it becomes robust with an increasing number of variants of each *N*-mutant (i.e. a variant with *N* amino acid substitutions). The average effect, 〈Δ*F*〉, may be biased by IFS (which are in principle infinite) and these are therefore not used in the fit of *F*_wt_ and 〈Δ*F*〉 in steps one and two (see Methods).

In the second step, effects of individual substitutions are estimated by an approach similar to that used in the first step (Fig. 1f). For each substitution, *s*, Δ*F_s_* is estimated from the subset of variants that contains that substitution. The effect is included in Eq. 2 by replacing one term of the average effect with the specific stability effect of *s* to be estimated with the remainder of the terms described by an average effect:

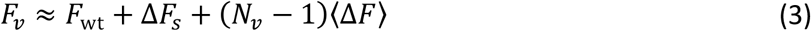

A robust fit to Eq. 3 requires a sufficient number of *N*-mutants (to estimate the average brightness) for several different values of *N*, but in contrast to above we only consider the subset of variants that contains the substitution *s*. To illustrate the required number of variants per substitution, Fig. 1f shows three examples of fits to Eq. 3 where it is clear that the 14 observed variants containing L178V result in average values of the *N*-mutant brightness that do not fit the model well. In contrast, the enhancing V163A and detrimental C48S substitutions, both fit the model well with 677 and 194 observed variants, respectively.

When all initial effects have been estimated by this mean-field approach, we perform a global optimization in step three using damped least-squares optimization from which uncertainties are calculated (see Methods).

The three-step estimation strategy ensures that effects that are poorly estimated due to few observations, typically < 10-20 depending on the distribution on *N*-mutants (supplementary Fig. S1), do not affect the initial estimation for substitutions with good data. A 5-fold cross validation shows that the model is able to predict the brightness of GFP variants not used in training accurately with an RMSD on brightness of 0.25 (ca. 10% of the dynamic range from 1.4 to 3.8) and a Spearman’s correlation coefficient *r*_s_=0.87 (Fig. 2). Notably, the most substituted variants are predicted well with 91% of active variants (brightness >2.7) correctly predicted as active for variants with six to eight substitutions and 73% for variants with nine or more substitutions.

**Figure 2.**
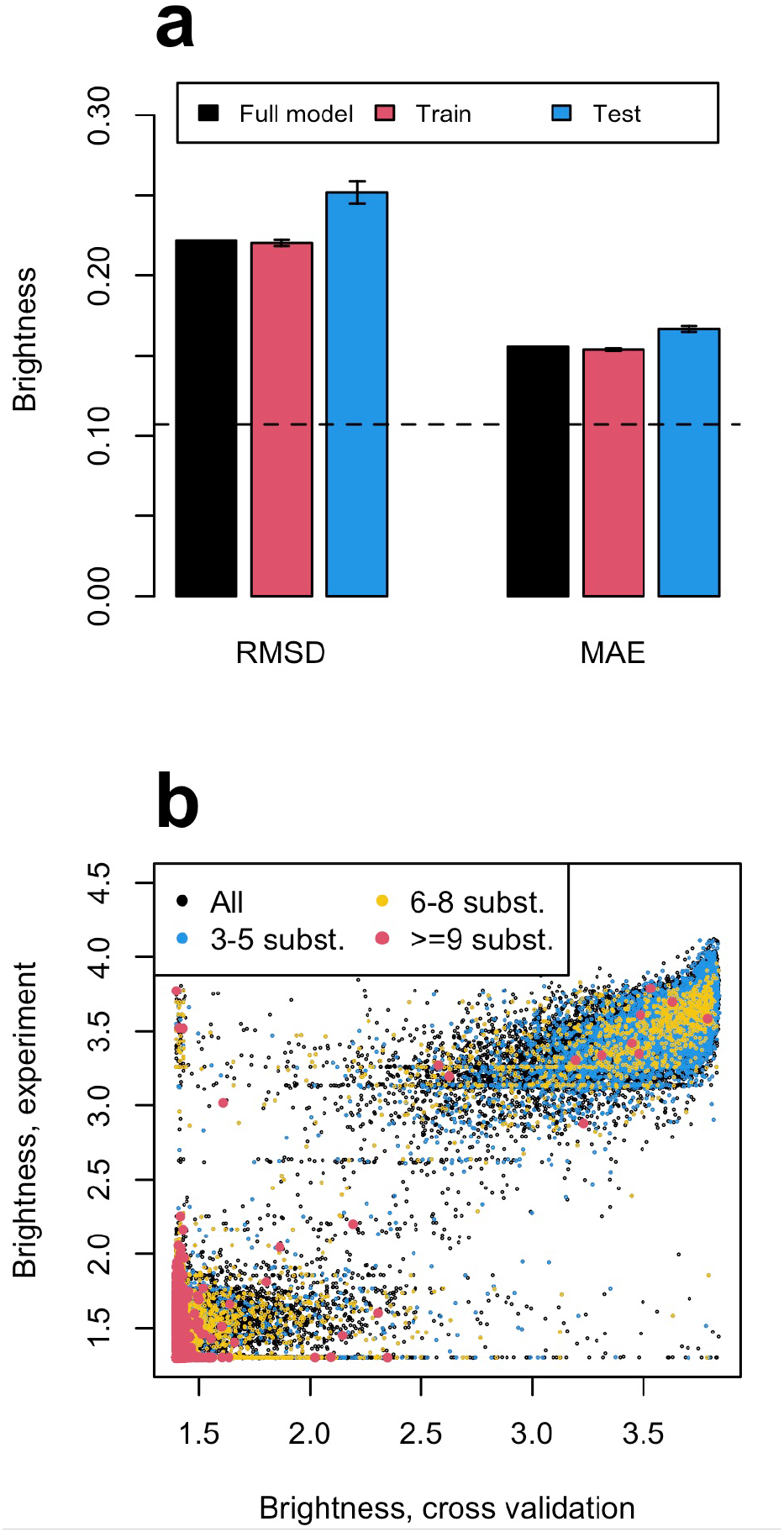
Five-fold cross validation of GFP multi-mutants where each variant was randomly assigned to one of five groups. (a) Root mean square deviation (RMSD) and mean absolute error (MAE) of predicted brightness for the full model fitted to all data (black) and the mean of the training and test data of the five cross validations (red and blue respectively). The error bars show the standard deviation from the five models each trained on ~80% of data and the dashed line indicates the experimental standard deviation of the wild-type brightness. (b) Predicted vs. experimental brightness of all variants, where the prediction of each variant is from the cross-validation when used as test data. The variants are colored according to the number of substitutions to show that even the mostly mutated variants are well predicted.

GMMA is based on a thermodynamic model and it is interesting to considered to which extent our fitness potential may relate to the thermodynamic stability as measured by chemical unfolding. We estimate the potential of the reference sequence (avGFP + F64L) to be *F*_wt_ = −2.8 using room temperature and a gas constant corresponding to units of kcal/mol. This is substantially smaller in magnitude than values < −10 kcal/mol reported from unfolding experiments of αGFP (avGFP + Q80R:F99S:M153T:V163A); however, unambiguously determining the stability of GFP is challenged by at least one folding intermediate of stability −3.7 kcal/mol [20]. The comparison is further complicated by *in vitro* unfolding experiments being qualitatively different from the equilibrium probed by GMMA. As discussed above, GMMA may probe the stability of the pre-mature of GFP and its ability to covalently and irreversibly mature the fluorophore in the context of a cell, whereas unfolding experiments deactivates the mature protein *in vitro*. Since maturation is spontaneous, the pre-mature state is likely less stable compared to the mature protein and thus, GMMA could probe a less stable structure compared to unfolding experiments of the mature protein.

We tested the robustness of the conclusions and values obtained by GMMA using synthetic data. Specifically, we generated multi-mutant data resembling those obtained for GFP and asked whether GMMA could recover the variant effects that were used to generate the data (see Methods). This analysis showed that it is possible to recover quantitatively the variant effects when the global model that relates the fitness potential to the experimental readout is accurately known. Importantly, however, the analysis also revealed that GMMA can robustly recover the ranking of the variant effects and distinguish those that are beneficial from those that are not even when the global model is imperfect (Supplementary Fig. S2). In that case, however, the scale of the estimated fitness potential might not be accurate.

### GFP substitution effects

Using GMMA we obtain accurate estimates for 1,107 substitution effects (59% of the 1,879 present in the library) with 80% found to decrease and 8% enhance the fitness potential, and the remainder to have an insignificant or neutral effect (Fig. 3). In the following we highlight some points that can be learned about GFP from these.

**Figure 3.**
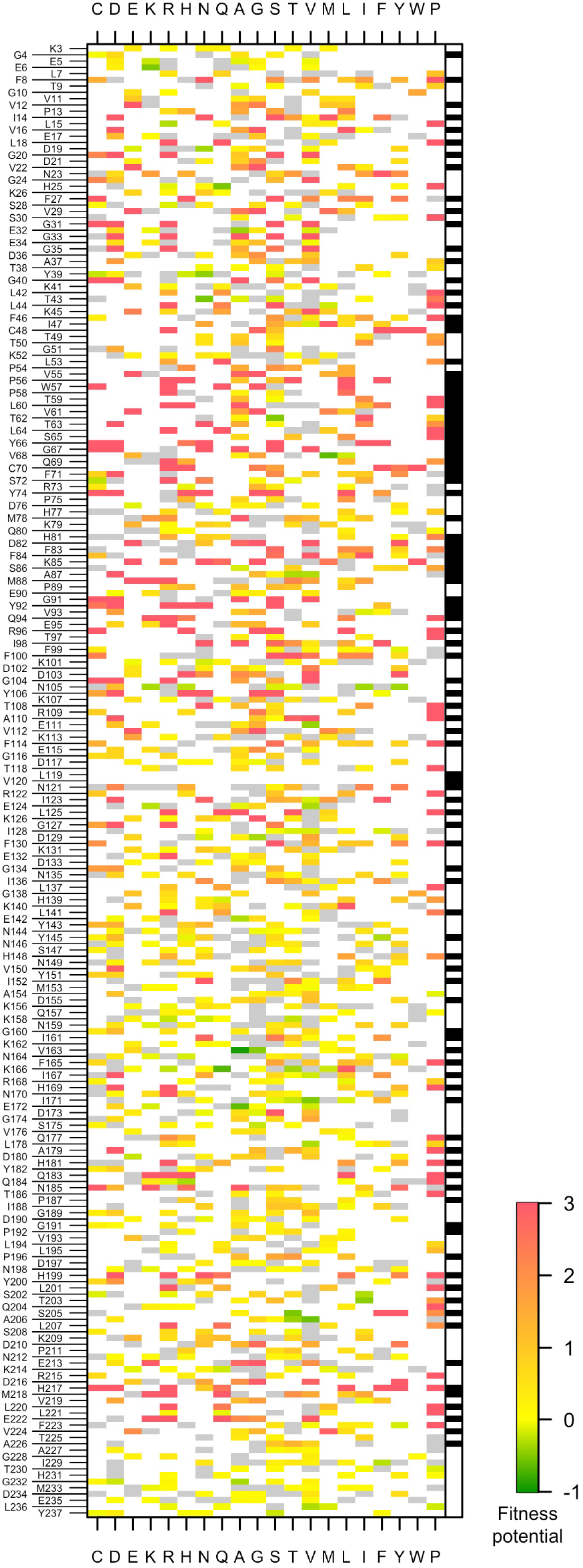
Heatmap showing the 1107 single-substitution effects estimated by GMMA from the multi-mutant library of GFP. Green indicates an enhancing substitution, yellow are substitutions with close-to-zero effect, and orange to red indicate levels of substitutions that diminish the fitness potential. Substitutions in gray are observed in the library but poorly estimated and white substitutions are not observed in the data. The rightmost column shows the solvent exposed residues in white and buried residues in black.

The majority of enhancing substitutions are found at solvent exposed positions (63/83; 76%; Fig. 3) together with almost all substitutions with insignificant effect, i.e. Δ*F_s_* close to zero (126/140; 90%). Both percentages are enrichments compared to the overall fraction of exposed substitutions (712/1,107; 64%). Nearly all substitutions at buried positions have detrimental effects (361/395; 91%) with the notable exception of the top-ranking substitution, V163A, which enhances the potential 38% more than the second best substitution, K166Q, and has low uncertainty due to a large amount of observed variant contexts in the data (supplementary Fig. S3). Another indication that V163A is highly beneficial is that another substitution at this position, V163G, is found to be the 12^th^ most enhancing substitution.

A group of positions with enrichment of enhancing substitutions is solvent exposed Phe, Ile and Leu for which 11/64 (17%) are stabilizing. With the exception of two Ile to Phe substitutions, these tend to slightly decrease hydrophobicity which is in apparent contrast to other engineering studies that find increased surface hydrophobicity to be stabilizing [21,22] but fits well with the observation that such stabilizations tend to have poor additive contributions [22].

Surface positions with wild-type Glu have a significantly higher fraction of enhancing substitutions, 12/71 (17%), as compared to Asp with only 5/90 (6%) substitutions being enhancing. This may suggest a qualitative difference in the role of these two negatively charged amino acids (that otherwise have the same transitions in the codon table) in the context of a beta-barrel. Furthermore, four of the 15 most enhancing substitutions remove a Glu sidechain from the surface (Fig. 4).

**Figure 4.**
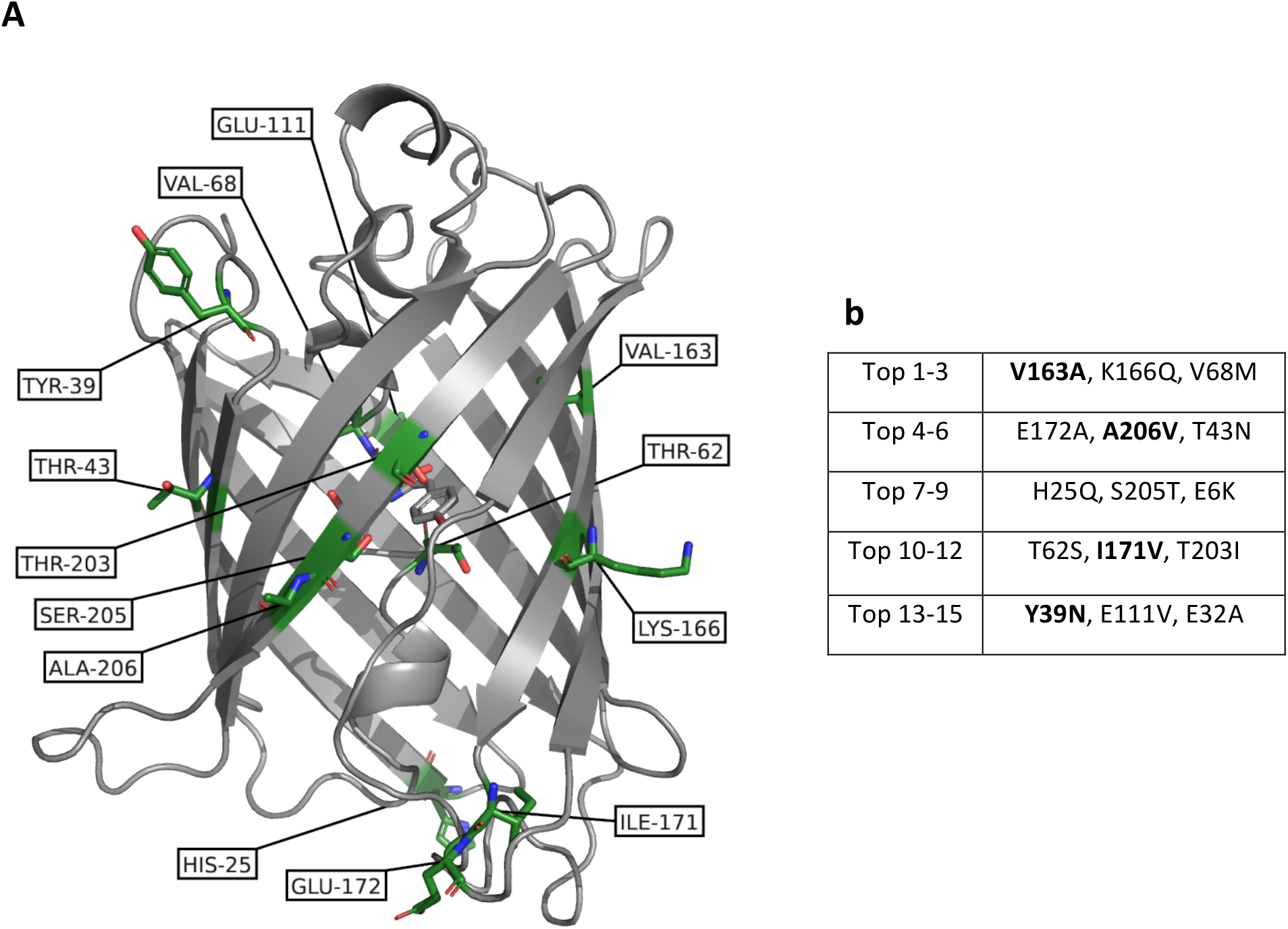
Positions (a) of the top 15 enhancing substitutions (b) shown on the structure of GFP (PDB entry 1EMM). Position 163 has two substitutions within top 15 and rank 9 is not resolved in this structure. Substitutions known from sfGFP are marked in bold in (b).

All 42 substitutions at all ten Pro residues in GFP are found to be detrimental, which is not surprising considering the structural role of Pro. Other positions that only show highly detrimental substitutions include the chromophore positions Y66 and G67 and the maturation-related sites R96 and E222 [23].

To examine whether the variant effects obtained from GMMA could be rationalized by structural stability of the mature state of GFP, we used Rosetta to predict the change in thermodynamic stability for all individual substitutions. Overall, we find a relatively good correlation between GMMA and Rosetta (*r*_s_=0.6; supplementary Fig S4a) suggesting that structural stability indeed is a substantial component of the fitness potential. However, when we focus on those substitutions identified by GMMA as most beneficial we find that most are not identified by the structural stability calculations since none of the top 15 substitutions overlap (supplementary Fig S4a). We further find that the Rosetta calculations correlate more strongly with the results of GMMA than of the measured brightness of singly-substituted variants (*r*_s_=0.6 vs *r*_s_=0.4, respectively; Fig. S3), lending further support to the accuracy of GMMA.

### Engineering GFP based on GMMA

Having used GMMA to identify individual substitutions that can suppress the detrimental effect of other substitutions, we examined whether these substitutions alone could be combined to improve GFP. Thus, we progressively inserted the 15 most enhancing substitutions into GFP in sets of three (Fig. 4b) and measured fluorescence levels during expression in *E. coli* at 41 °C. Measuring fluorescence directly in cells was intended to resemble the conditions of the assay used to generate the data [7] while the elevated temperature was applied to enhance the possible associated differences in thermal stability. As references we compare to the reference GFP sequence used in the screen (WT) and sfGFP (see Methods).

We find that all variants show fluorescence levels that are higher than the reference sequence (Fig. 5 and supplementary Fig. S5) with the Top6 variant giving as high an in-cell signal as sfGFP. Notably, Top6 and sfGFP only have two substitutions in common, V163A and A206V, (Fig. 4b and supplementary Table S1) whereas sfGFP contains 10 substitutions relative to WT. While the Top12 and Top15 variants have lower fluorescence than Top6 and 9 they are still at least as bright as the reference wild type. We expect the increased fluorescence mainly to reflect a difference in protein levels (i.e. not increased intrinsic fluorescence) and thus a proxy for *in vivo* maturation and stability.

**Figure 5.**
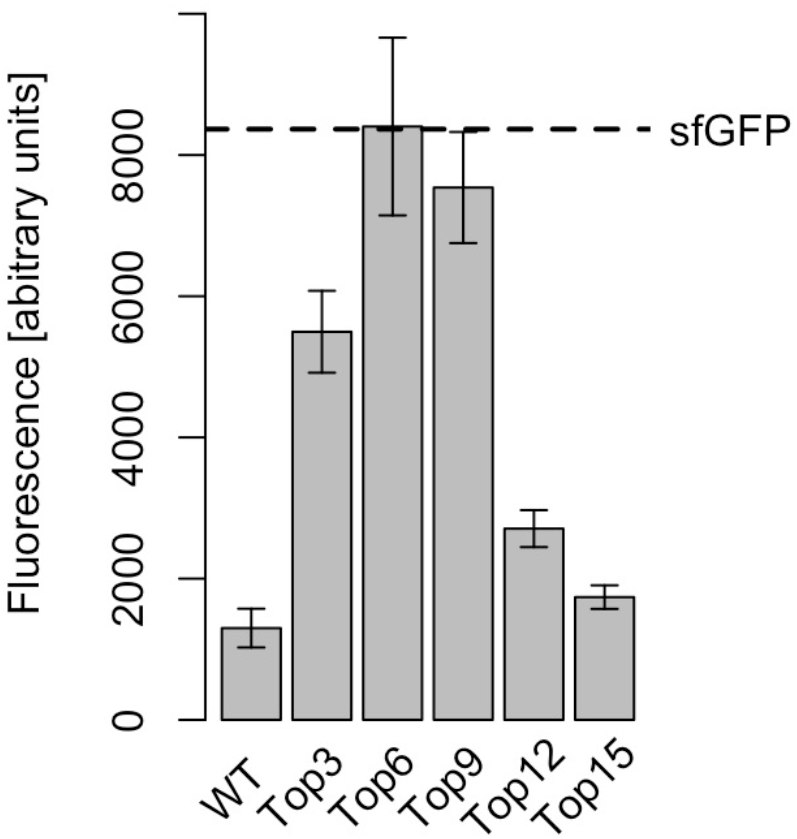
Experimental test of multiply-substituted GFP variants composed of top ranking substitutions identified by GMMA. Error bars show the standard deviation between four replicates.

These experiments demonstrate two important points of GMMA. First, that at least the six top-ranking substitutions appear to have a substantial additive positive effect, with Top6 giving significantly more signal than Top 3. One might speculate that the leveling off of Top9 and the decreased signal of Top12 and Top15 relative to Top6 be caused by context-specific effects, due e.g. to the structural proximity between I171V and E172A or T203I, S205T and A206V (Fig. 4). The latter combination, however, appears to be close to optimal in other high-throughput experiments [24]. Second, none of the 15 most highly ranking substitutions are highly deleterious (all variants are fully functional). This observation supports the robustness of the outcome of GMMA since these enhancing substitutions have all been observed in a diverse set of multi-mutants in which they have been judged to play an enhancing role.

In addition to the calculated uncertainties, GMMA offers transparent analysis of the final fits of individual substitutions (supplementary Fig. S2). These reveal that effects of the top 5 ranking substitutions are well-determined by the data, whereas rank 6 (T43N) is only observed in nine variants and thus we have less confidence in this. While V163A and A206V are both present in sfGFP, the high fluorescence of Top6 suggests that K166Q, E172A and V68M may be promising substitutions for GFP engineering. Continued experimental investigation of these specific effects is interesting, but beyond the scope of the present study.

### Known GFP substitutions

To examine whether the substitutions we identify using GMMA more generally have enhancing and background-insensitive effects, we compare these to substitutions that are known to be beneficial for GFP in multiply substituted contexts found in the literature [18,19,25—28] (see Methods). Known substitutions are in general found to be enhancing or with insignificant effect (Fig. 6a) and GMMA outperforms both Rosetta stability calculations and experimental measurements of single-substitution effects in the identification of known substitutions (Fig. 6b). Known substitutions that are estimated by GMMA to have a detrimental effect (supplementary Table S1) may indeed also be accurate because they were originally selected for other purposes. Perhaps most notable are the chromophore substitution S65T, selected for spectral properties, and C48S, introduced to avoid cysteine oxidation in extracellular environments. Likewise, the substitutions Q80R and H231L, here estimated to be slightly detrimental, are historical substitutions caused by early PCR errors which are still present in some synthetic genes of GFP variants [29].

**Figure 6.**
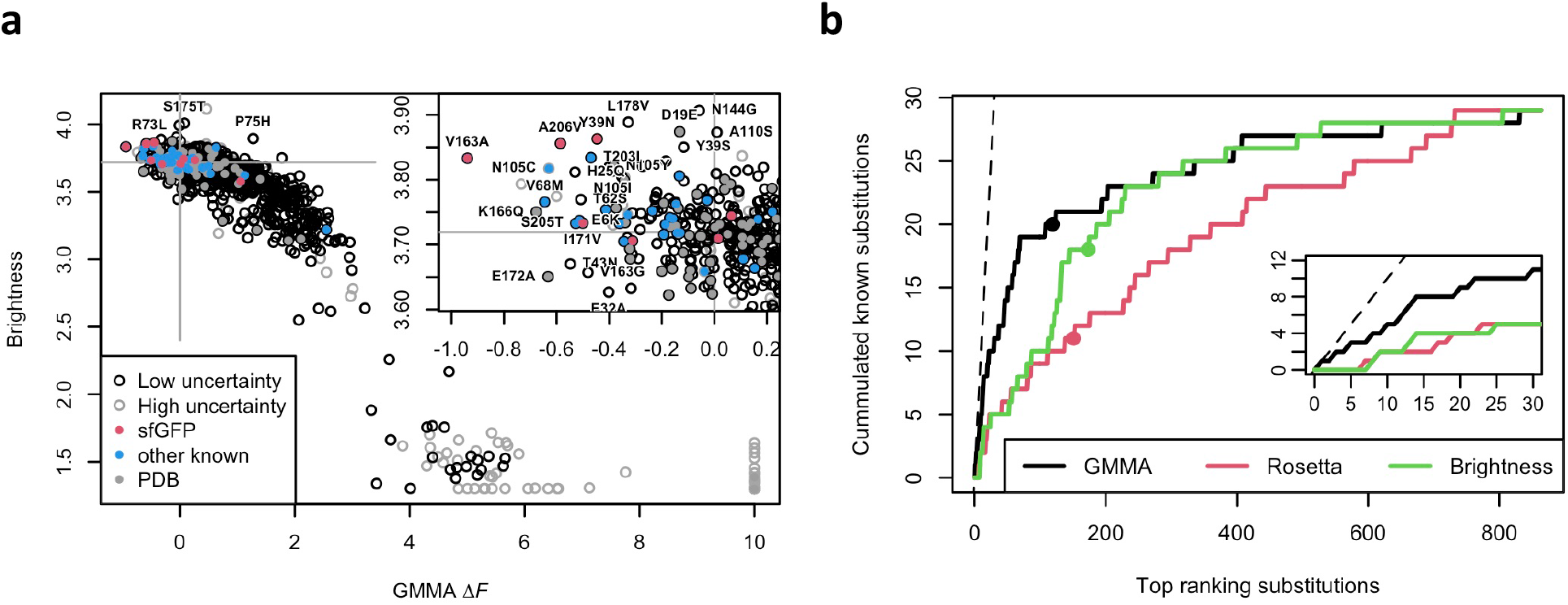
a) Brightness versus GMMA stabilities for single amino acid variants. GMMA estimates with large uncertainties are shown in grey circles. Substitutions known from optimized versions of GFP are shown in color. The vertical lines mark Δ*F*=0 and the horizontal lines the brightness of the wild-type sequence. The insert shows the highly enhancing, high brightness region and is enriched in known GFP variants. b) The graph shows the number of known substitutions within a given top-ranking number of substitutions according to GMMA (black), Rosetta (red) or singly-substituted variant fluorescence measured in the original data (green). The plot considers 29 unique known substitutions among the 863 low uncertainty substitutions for which single-variant fluorescence measurements and Rosetta calculations were available (see methods). The points mark the substitution with effect closest to zero, i.e. closest to wild-type, for each measure (gray lines in panel a). The dashed line indicates maximum performance, i.e. the 29 known substitutions as the top ranking 29 substitutions.

We furthermore consider 136 substitutions found in GFP variants deposited to the PDB. As expected, these are found to be enhancing or with insignificant effect. Of the 136, only four substitutions were found to have a slightly detrimental effect (Δ*F* < |*F*_wt_|/2) which confirms that GMMA robustly avoids deleterious substitutions also on a larger set of substitutions. In total, we find that 19 of the top 30 enhancing substitutions obtained from GMMA are known and that none of the known substitutions are estimated to inactivate GFP (Δ*F* > |*F*_wt_|).

The effects estimated by GMMA correlate strongly with the observed brightness of singly-substituted variants (Fig. 6a; *r*_s_=-0.77). This, however, is mainly caused by the fact that the averaged FACS readout and GMMA analysis agree on low brightness variants. In contrast, when the goal is the more challenging one of identifying enhancing variants we see a much weaker correlation between the observed brightness and GMMA (*r*_s_=-0.26 for variants identified by GMMA as significantly enhancing; see Methods) because individual enhancing variants have only modest effects on brightness when introduced in an already stable background (Fig. 6a, insert).

## Conclusions

We have here introduced a concept to quantify functionally-enhancing effects of individual substitutions through a global analysis of high-throughput assay measurements of multi-mutants. We demonstrate that the effects of single amino acid substitutions may be better determined by analysis of multi-mutants and have presented a Global Multi-Mutant Analysis (GMMA) that implements this.

Our fitness potential is shown to correlate with structural stability (Fig. S4), with the important addition, e.g. compared to *in vitro* unfolding experiments, that substitutions that decrease function appear as destabilizing. Our model is based on thermodynamics, Eq. 4, and the fitness potential may be interpreted as a free energy (in kcal/mol) under the given assumptions; most notably the absence of couplings between substitutions and the global map between the fitness potential to the assay readout. Experiments on synthetic data furthermore indicate that the ranking of substitutions and zero-point (fraction of enhancing substitutions) are robust towards inaccuracies in the global model and thus, that GMMA can be interpreted as a free energy with a system dependent scale.

A cross-validation analysis demonstrates that GMMA can be used to predict experimental results on a complex fitness landscape involving nine or more substitutions in many variants which is also confirmed experimentally by fluorescence measurements of multiply-substituted GFP variant that contains the top ranking GMMA substitutions. We show that the six best substitutions may be combined to design a GFP variant which, under the assay conditions, is highly fluorescent and identify at least three enhancing substitutions, K166Q, V68M and E172A, that are known but seems underestimated in the literature of enhanced GFP variants. Finally, we show that GMMA is able to identify a large number of the substitutions that are known to be beneficial to GFP in multiple-substituted contexts.

## Data and code availability

The code and dataset generated and analyzed during the current study are available in the GitHub repository, https://github.com/KULL-Centre/papers/tree/master/2020/multi-mutant-analysis-Johansson-et-al.

## Acknowledgements

We thank Prof. Amelie Stein for discussions and insights and Charlotte O’Shea for expression and fluorescence measurements of the GFP variants reported here. This work was supported by Independent Research Fund Denmark and the PRISM (Protein Interactions and Stability in Medicine and Genomics) centre funded by the Novo Nordisk Foundation (NNF18OC0033950).

## Contributions

KEJ, KLL and JRW conceptualized the method. KEJ, KLL and JRW developed the method and analyzed the data. KEJ implemented the software and wrote the manuscript with contributions from all authors.

## Methods

We explore a thermodynamic model and assume that the observed brightness, *B*, of a variant, *v*, is proportional to the fraction of the protein found in the active state A:

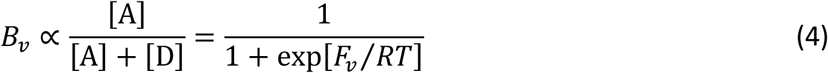

where [A] and [D] are the concentrations of active and inactive protein respectively, *R* is the gas constant and *T* the temperature. Following the original report of the data [7], the log fluorescence, or brightness *B*, is assumed proportional to the fraction of active protein. If the proportionality of the brightness and fraction of active protein is an accurate description, the fitness potential may be interpreted as a free energy that describes the equilibrium *A* ⇌ *D*. Throughout this work, we use a value for *RT* that corresponds to a fitness potential in kcal/mol at room temperature. We carried out all data analyses using the *R project for statistical computing* with packages *minpack.lm* and *igraph*.

### Initial estimation

The initial estimation of stability effects considers the *average* effect of substitutions, 〈Δ*F*〉, which is sensitive to highly destabilizing substitutions. Thus, we first detect *irreversible fatal substitutions* (IFS) that are inactive in all contexts and have an unlikely pattern of activity among multi-mutants, e.g. many inactive double mutants in the case of GFP (see supplementary Appendix 1). We stress that the IFS are detected directly from an analysis of the data and is not based on any pre-defined notions of variants or regions that are particularly relevant for function. Variants containing nonsense mutations are also generally expected to be IFS. We exclude 62 nonsense IFS and 51 missense IFS from the initial estimation together with the 2,310+4,851 variants that contain these nonsense and missense IFS substitutions respectively.

The wild-type fitness potential, *F*_wt_, is first estimated together with the average effect of all substitutions, 〈Δ*F*〉, using Eq. 5 which combines Eqs. 2 and 4. These are fitted to the average brightness of the *N*-mutants, 〈*B*〉_*N*_, i.e. the average brightness of double-mutants, triple-mutants, etc.

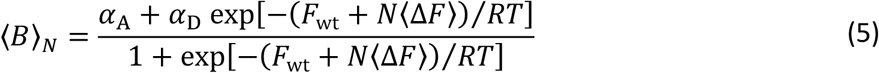

Here, a constant baseline, *α*_D_, for the brightness of the inactive variants is fitted whereas a constant baseline for active variants, *α*_A_, is not fitted independently but calculated from *F*_wt_ and the brightness of the wild-type sequence, *B*_wt_, during fitting:

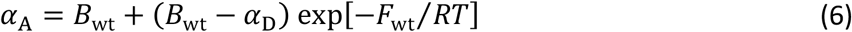

This makes *α*_A_ less sensitive to noise and outliers in the variant readout by relying on the data point (*F*_wt_, *B*_wt_) which is experimentally well-determined with 3,645 barcodes in the high-throughput assay, i.e. individual observations of wild-type nucleotide sequence (2,444) or synonymous sequences [7]. A standard error of each parameter is calculated as the square root of the diagonal of the inverse Hessian matrix. The initial wild-type potential is estimated to *F*_wt_ = −1.8 ± 0.2, the average effect of substitutions 〈Δ*F*〉 = 0.45 ± 0.04, and *α*_D_ = 1.43 ± 0.03 (Fig. 1e). This results in *α*_A_ = 3.8.

In step 2, we use the values of *F*_wt_, 〈Δ*F*〉 and the baseline parameters from step 1, to estimate initial values of the individual substitutions, Δ*F_s_*, from the subset of variants that contains the substitution, *s*, using Eqs. 3 and 5 (Fig. 1f). The use of a fixed value of 〈Δ*F*〉 makes the initial estimates robust and self-consistent for the global fit, and the approach is further supported by the observation that variant effects are mostly independent of the details of the background [15]. Since we always only change the background with a single substitution, 〈Δ*F*〉 is not expected to vary much and in practice less than the uncertainty that results from small sets of variants. We require that all subsets have a diversity of at least 3 different multi-mutants that spans the transition region and gives a fit with standard error < 1.0 and an absolute deviation < 5.0 log fluorescence units. With this, we find that 56% of substitutions have sufficient data for this initial estimation. We then afterwards estimate the remaining initial Δ*F_s_* values from these well-determined effects.

### Graph analysis

We pre-process the data for the global multi-mutant analysis and build a bipartite graph by assigning protein variants to one layer of nodes and another layer of individual substitutions. All variant nodes are linked to the substitution nodes that the variant is composed of. We check that all nodes are connected in the graph. If a subset of variants is composed of a subset of substitutions that does not occur in the rest of the variants, this graph becomes disconnected from the rest, and GMMA may be carried out on the subset alone (e.g. a single-mutant library is fully disconnected, GMMA cannot be applied to this). Single mutants do not inform the global fit more than any other variants and indeed 130 stability effects are estimated without the single mutant being observed. Substitution nodes with a single link represents substitutions that contributes one parameter and only one data point to the global analysis, referred to as hanging nodes. These do not inform the global optimization and, thus, the effect of these are calculated after the global optimization. In summary, the graph is cleaned for 7 disconnected node pairs, and 257 hanging substitution nodes together with 255 dependent variants nodes. Finally, the graph is examined to ensure that no pair of substitutions only occurs together as these would make them impossible to distinguish and should be reparametrized as a single effect. The graph analysis is independent of the initial estimates and considers all data, including IFS and non-sense substitutions.

### Global estimation

With input from both the initial estimation and the graph analysis, we perform the global optimization. The global fit was not stable without some type of regularization (presumably because of parameter correlations) and thus, we apply a trust region as implemented in the Levenberg-Marquardt least-squares algorithm with effects limited to the range −5 to 10. This is chosen over an elastic net type of regularization because we are interested in the fitted parameter values and these are not expected to follow a single normal distribution (e.g. centered at zero) and thus, a more complex prior would be needed, e.g. for ridge regularization. We optimize all effects {ΔF_s_} and the wild-type fitness potential *F*_wt_ to the nearest optimum (Fig. 1g). The baselines are fixed at the values determined in the initial fit (1.4 and 3.8). With analytically calculated gradients, the global optimization of 1,616 parameters from 53,763 data points took 4-5 hours on a normal laptop.

The fit has a reduced chi-square 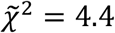 which indicates that some parts of the data do not fit the model; however, we note that the cross-validation analysis shows that the data is robustly fitted and with at most a small level of overfitting (Fig. 2). Our fit appears notably better than that reported previously for similar models [6,7] which we expect to be related to our inclusion of the global parameter, *F*_wt_. One contribution to an elevated 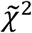 could be the use of the distribution of the observed brightness of the wild-type sequence as a proxy for the uncertainties of all variants, which may indeed have higher uncertainties in the assay.

### Uncertainties

In the global analysis, we estimate the uncertainties from the covariations in the inverted Hessian matrix. We calculate two error measures to judge the uncertainty and reliability of the effects. The first, *δ_s_*, is calculated using the log-fluorescence measurement uncertainty, reported to be *δB*_wt_ = 0.11 for the wild-type sequence [7], propagated via the covariance matrix diagonal and multiplied by 3 to get the 99.7% percentile (and to somewhat compensate for the expectation that the variants have higher uncertainty than the wild type):

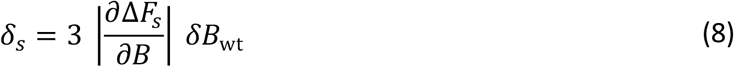

A resampling experiment suggests that this measure captures the estimation accuracy well (supplementary Fig. S1).

The second uncertainty, *δ*_Δ*Fs*_, is used to filter out unreliable stability effects and is calculated from:

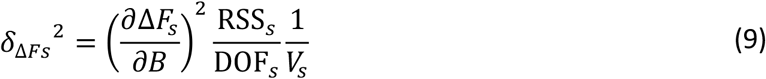

Again, the derivative is from the diagonal of the covariance matrix, RSS_*s*_ is the residual sum-of-squares of the variants used to estimate substitution *s*, DOF_*S*_ is the number of degrees-of-freedom and *V_s_* is the number of variants used to estimate substitution *s*. The last factor gives an error-of-the-mean type of uncertainty that compensates for the case where a lucky fit of few variants is penalized. The number of degrees of freedom for a substitution assumes that a uniform fraction of parameters is estimated together with Δ*F_s_* from the *V_s_* data points

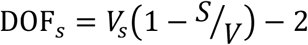

where *S* and *V* are the total number of substitutions and variants respectively. We set a relatively conservative threshold and mark 772 (61%) effects with *δ*_Δ*Fs*_ > 0.05 as unreliably (a value that can be compared to 〈Δ*F*〉 ≈ 0.5). This low threshold has been set by manual inspection of plots that show the fit of each substitution to its respective subset of variants (similar to Fig. 1f). We use a constant threshold here to facilitate a clear discussion. However, in a specific application, substitutions could be judged individually based on both uncertainty measures and such plots, since many of the 772 poorly estimated substitutions are still informative (as in supplementary Fig. S3). Notably, 222 (12% of all substitutions) are exclusively or predominantly observed in inactive variants and are therefore only represented in the flat region of the model. Thus, all of these may reliably be identified as destabilizing or even well determined on a range, even though the reported point estimate itself is highly uncertain. Of the remaining 550 (29%) substitutions with uncertain effects, the majority (398, 21%) are caused by poor statistics with five or fewer observed variants. The conservative threshold does exclude some substitutions with more than 10 observations (61 or 3%), that are potentially stabilizing, e.g. L221V, Q80K or E6A, and may be interesting depending on the application.

### Synthetic data

Each of the 1615 substitutions considered in the global analysis (i.e. those not disconnected or hanging) are assigned their estimated effects if these pass the uncertainty threshold, and otherwise a random value drawn from a normal distribution resembling the accurate effects. Synthetic brightness values for 53,762 variants are calculated via Eq. 1, the value of *F*_wt_ given above and one of three global models relating brightness to fitness potential: 1) the sigmoid function as used in the global fit with the parameters given above (Fig. S2A). 2) a log-linear function *B_v_*=log(−10 * *F_v_* + 20) for *F_v_* < 0 and *B_v_*=log(5) otherwise (Fig. S2B). 3) a discontinuous, close to binary, log-linear function *B_v_*=log(—2 * *F_v_* + 30) for *F_v_* < 0 and *B_v_*=log(5) otherwise (Fig. S2C). The synthetic brightness is added a Cauchy distributed noise of scale 0.05 (compared to baselines 1.4 and 3.8) and explicit misclassification of 158 random variants (similarly to estimated misclassification rate in the GFP data). Since the Cauchy distribution is not unlikely to generate samples of very high absolute values, brightness values above 4.5 are returned to the model value and values below 1.28 are set to 1.28 (minimum brightness observed in GFP data). GMMA of the three synthetic data sets is carried out as described above except that the initial mean-field estimate is omitted and random initial values are used instead.

### Classification

For the sake of discussion and early IFS identification, we classify all variants as either active or inactive. Variants with brightness below 2.7, half-way between maximum and minimum observed brightness in the original data, are assigned as inactive and the rest as active.

We mark substitutions with low uncertainty in the fitness potential from GMMA (high/low uncertainty classification described above with the uncertainty calculation) as *significantly enhancing* if the effect plus uncertainty is less than zero, *significantly detrimental* if the effect minus uncertainty is greater than zero and *insignificant* otherwise. Substitutions with high uncertainty are marked as detrimental if the effect is greater than the wild-type potential and marked as unknown otherwise.

Solvent exposure categories are exposed or buried according to DSSP [30].

### Rosetta stability effects

Rosetta scores were calculated according to protocols published in [31,32] using precompiled Rosetta version 3.13 of June 2021 with the REF15 score function (ref2015_cart) and the chromophore parameters distributed with Rosetta (CRO.params). Calculations were based on PDB entry 1EMM that covers position Leu-5 to Ile-227 both included. This was prepared by 20 independent and unrestrained minimizations using the command:

./relax -score:weights ref2015_cart -relax:script cart2.script -use_input_sc -relax:cartesian -ignore_unrecognized_res -relax:min_type lbfgs_armijo_nonmonotone -fa_max_dis 9.0 -extra_res_fa CRO.params

The lowest energy structure was used to evaluate all single variants using the command:

./cartesian_ddg -score:weights ref2015_cart -ddg:iterations 3 -force_iterations false -ddg::score_cutoff 1.0 -bbnbrs 1 -fa_max_dis 9.0 -ddg::cartesian -ddg::legacy false -extra_res_fa CRO.params

We used no special handling of substitutions involving proline and the substitution effect was calculated as the average of three repeats of the calculation. Results in Rosetta energy units (REU) were divided by 2.94 which have previously showed to correlate well with stability effects in kcal/mol [31]. Nonsense and chromophore mutations could not be evaluated and calculations at position Arg-80 were ignored because of the mismatch to our reference Gln-80.

### Known substitutions

We focus on published variants that are relevant for our reference sequence (avGFP + F64L) and consider the substitutions that constitute sfGFP [18], T-Sapphire GFP (tsGFP) [26], a split GFP (splitGFP) [27], a computationally-optimized GFP known as des11 [19] and a tryptophan chromophore variant called nowGFP [28] (supplementary Table S1). While many more variants exist in the literature, these five variants represent close to all substitutions [25]. We expect substitutions from these to be stabilizing or have an insignificant effect in GMMA. We note that some substitutions from nowGFP have been reported to support the tryptophan chromophore [33, 34].

Additionally, we also consider 136 substitutions found in 147 structures of GFP, selected in the PDB to have >90% identity to our wild-type sequence. Since these variants have all been expressed and crystalized, we expect that these substitutions do not destabilize GFP substantially, i.e. lower expectations compared to substitutions from the enhanced variants mentioned above.

### Experimental testing

The chromophore substitution S65T is known to change the spectral properties of GFP and since this is contained in sfGFP, we add this substitution to all variants (including the WT reference), in order to compare fluorescence levels measured using the same excitation wavelength of 488 nm.

Synthetic genes encoding the multiply mutated GFP variants were custom synthesized by Twist Bioscience in a pET29b expression vector and expressed using BL21(DE3) as host. The DNA sequence was codon optimized for *E. coli* for the reference sequence, and only varied at the positions of substitutions, where most common codons were used. Precultures were grown over-night and normalized for cell density to approximately OD_600_ of 3.0, respectively. Expression was carried out in 96-well plates which were inoculated with 5 μL in 45 μL LB medium containing kanamycin for plasmid selection. After incubation at 37°C for two hours, IPTG was added to 1 mM and the temperature was shifted to 41°C and fluorescence measured using a TECAN Infinite 200 Pro microplate reader over the next 12 hours. All measurements were done with two independent precultures, each with two technical replicates. The numbers reported in Fig. 5 are averaged over the two last hours of the profiles in supplementary Fig. S5.

