## Supplementary text and figures for "Global analysis of multi-mutants to improve protein function"

#### Supplementary figures and tables

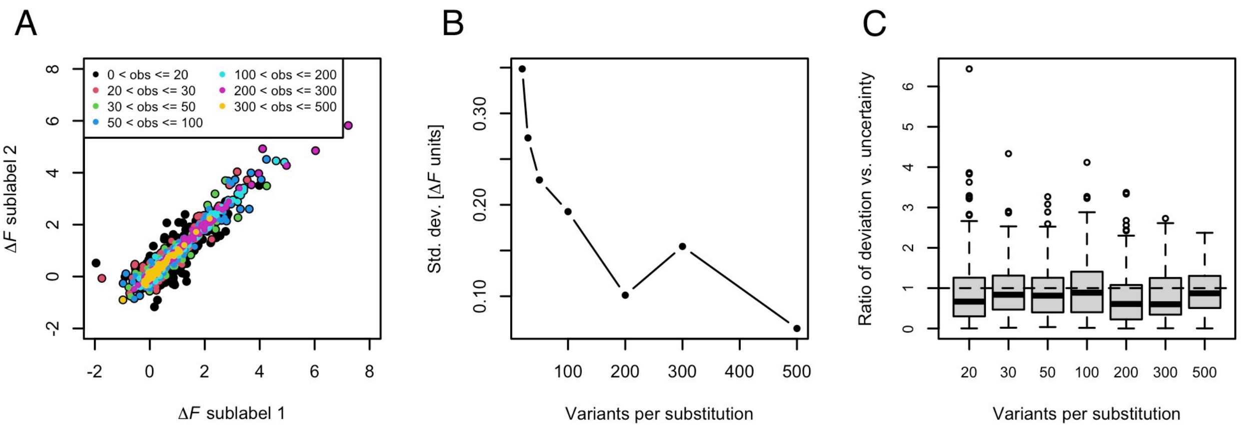


**Figure S1**. Validation of errors by sampling. We used the fact that substitutions are found in multiple variants to estimate the uncertainties of the variant effects. Each substitution was thus synthetically relabeled into two “sub-labels” (1 and 2) and these were assigned randomly to the different variants that contain this substitution, followed by a complete rerun of GMMA. Substitutions for which both sub-labels have uncertainty above the threshold are not included. (**A**) Correlation between effects estimated for sub-labels of the same physical substitution. Points are colored according to the number of variants that sub-label 1 was observed in. (**B**) Standard deviation from equal sub-label effects (spread around the diagonal in panel A) for the same bins as in panel A. Points are placed at the upper bound of the bin. As expected, when a substitution is found in many variants we can estimate its effect more precisely. (**C**) Boxplot of the ratio between the error-of-the-mean from sub-labels and the uncertainty, $\delta_s$ (Eq. 8) averaged for the two sub-labels. The box surrounds the two center quantiles and points outside an approximate 95% confidence interval are plotted individually. As a validation of our error estimate, a ratio of 1 is indicated with dashed lines to highlight that the reproduction of effects between sub-labeled substitutions is similar to the given uncertainties and in many cases smaller.

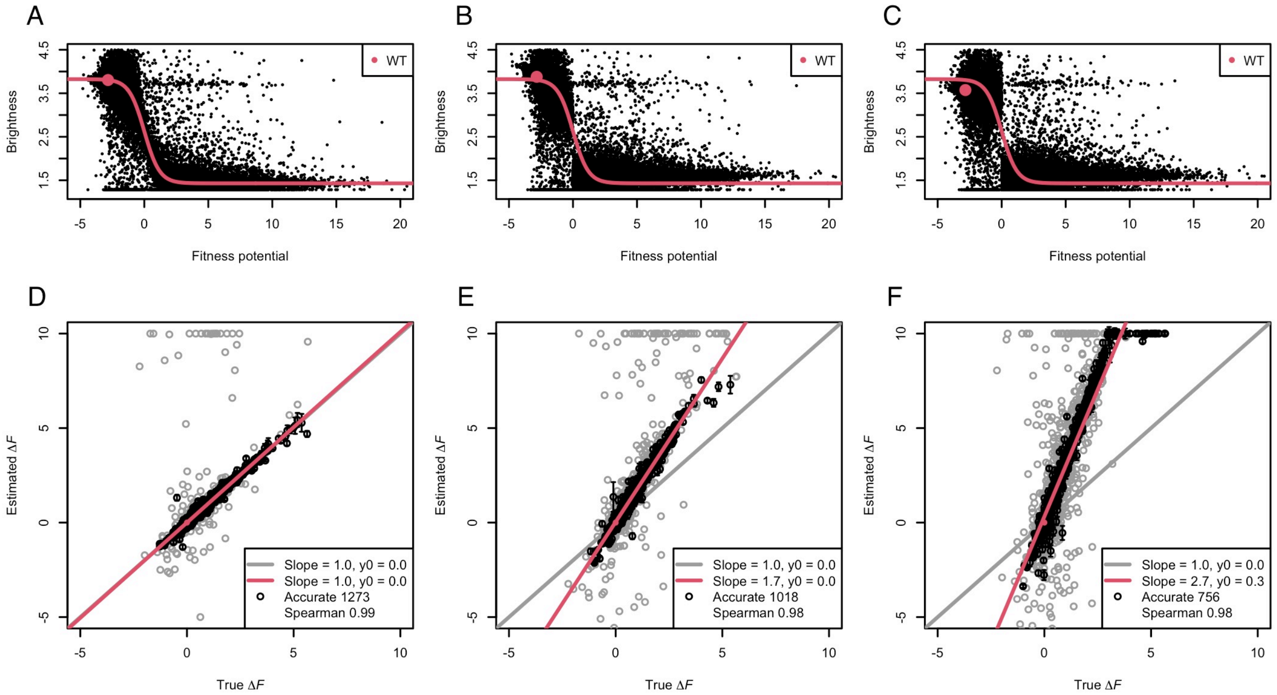


**Figure S2**. Three sets of synthetic variants (panels A-C) resembling the GFP data and the ability of GMMA to retrieve the true substitution effects (panels D-F). Using the same variants (i.e. combinations of substitutions) as for the GFP data and the estimated substitution effects, we generate synthetic fluorescence values using three global models designed to represent increasing levels of inaccuracy in the global function that maps the fitness potential to the assay readout (see methods): One identical to the sigmoid used in the global fit (panel A), a log-linear function (panel B) and a more dis-continuous log-linear model (panel C). All data are added substantial amounts of noise (see methods). For reference, panels A-C are all overlaid with the fitted model of the GFP data (red line). The ranking of the know substitutions effects are in all cases retrieved well with Spearman correlations of 0.99, 0.98 and 0.98. Likewise, the point of zero potential is also retrieved well, i.e. the fraction of enhancing and abating substitutions is accurately estimated with linear intersection coefficients of 0.0, 0.0 and 0.3 for the three global functions. The scale of the effects (linear slope) on the other hand, is only accurately retrieved when the global function used in the fit is an accurate description of the global function that generated the data.

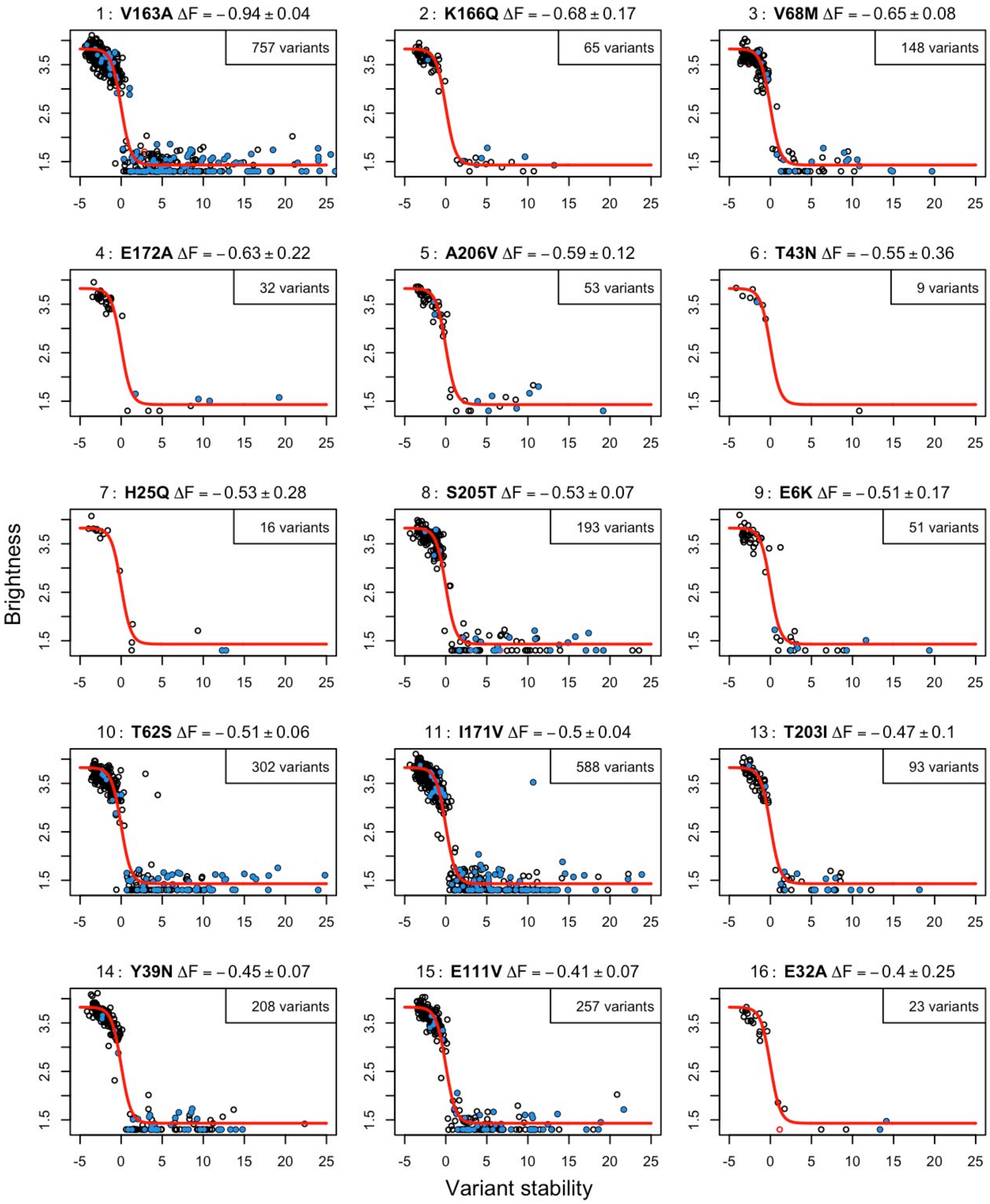


**Figure S3.** Final fit of the top 15 substitutions. Rank 12, V163G, is omitted because it overlaps with rank 1. All variants carrying the indicated substitution are plotted (black circle) with summed GMMA stability effect (x-axis; Eq. 1) and brightness (y-axis) together with the global model (red line; Eq. 5). Variants carrying more than 6 substitutions are marked (blue point). The few variants that are discarded based on the networks analysis (red circles) are not included in the indicated number of variants. Estimated stability effects are given after the rank and name of the substitution.

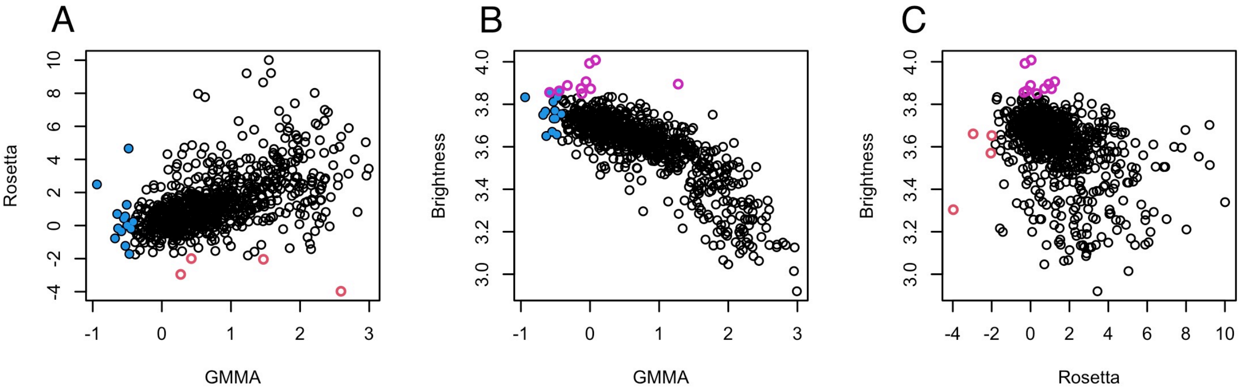


**Figure S4.** Comparing different measures of single-variant effects. The Spearman rank correlations between GMMA, Rosetta and brightness of single-substituted variants are 0.57, -0.77 and -0.43 for panels A, B and C respectively. All plots show the 840 substitutions for which all three measures were available and were measured to be active in the original data [7]. These include 14 of the GMMA top 15 (blue points), 4 of the Rosetta top 15 most stabilizing substitutions (red circles) and 10 of the top 15 brightest single-substituted variants (purple circles). Of all top-ranking substitutions only two overlap: A206V and Y39N which are both in the top 15 of GMMA and single-variant brightness. Including the 23 inactive variants results in slightly higher Spearman correlations of 0.60, -0.79 and -0.47 because these populate separate regions. One reason (among others) to exclude inactive variants here is that Rosetta is not expected to recognize substitutions that are deleterious to function but not structural stability, e.g. chromophore substitutions.

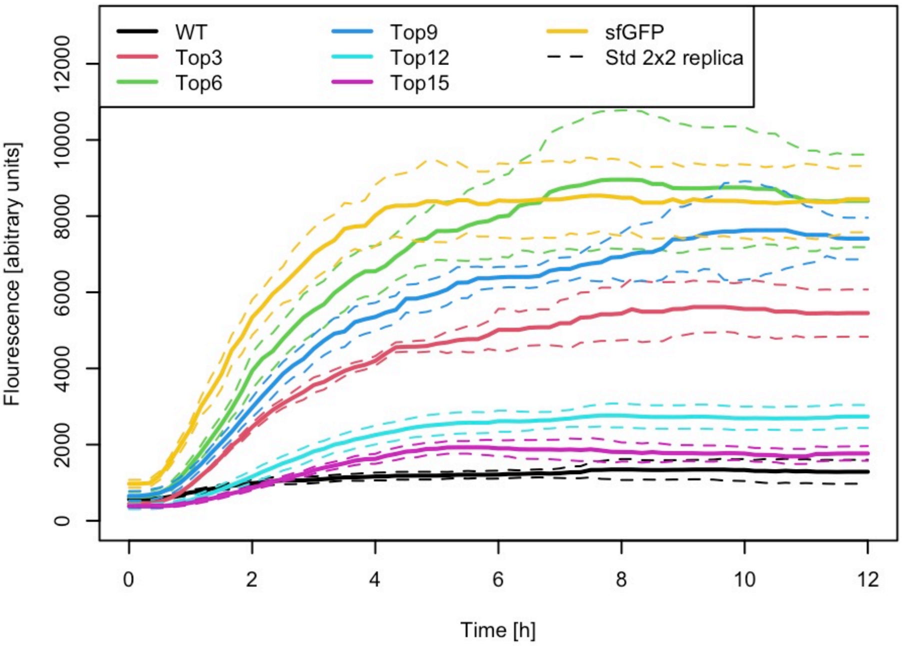


**Figure S5**. Fluorescence profiles of the tested GFP variants. Fluorescence is measured for 12 hours after induction of expression. All measurements are done in four replicates and the standard deviation is shown as dashed lines for each variant.

**Table S1**. Known substitutions

| | Stabilizing | Insignificant effect (neutral) | Destabilizing | Uncertain | Not in our data |
|---|---|---|---|---|---|
| sfGFP (10 of 11) | V163A, A206V, I171V, Y39N, Y145F | N105T, M153T | **Q80R,** F99S, **S65T** | | S30R |
| tsGFP (5 of 9) | V163A, T203I, Y145F, S175G | | **H231L** | | L62F, Q69M, C70V, S72A |
| splitGFP (17 of 21) | V163A, A206V, S205T, I171V, E111V, K166T, N105K, Y145F, I167V, F223Y, I128T | M153T | **Q80R**, K3Q, F99S, **C48S** | T225N | S30R, Y39I, C70A, L221H |
| des11 (7 of 13) | Y145F | K158N, A154P | **H231L**, N198Y, **S65T*** | N105C | S30R, K126H, S202H, Q204W, A206E, T225E |
| nowGFP (20 of 26) | V163A, V68M, E6K, I171V, K166T, I128V, T43S, N164Y | T230P, Q177L, M153T | **H231L**, N170D, Y151N**, N146I**, V150A**, **S65T**, L207Q** | L42M, M233A | V61K, Y66W, S72A, Y145A, H148G, A206K |
| PDB (136 of 220) | V163A, K166Q, E172A, A206V, S205T, I171V, Y39N, N105Y, D19N, G232A, E124V, L236V, A87V, Y145F, S175G, V68L, Y39H, I167V, E32V, I128T, N105S, S202C, V93I, N164Y, Y145H, H25R, K101E, T9A | Y237S, K101N, N164H, L236M, D19E, N105T, E5A, E124A, D190N, V219I, K166G, M153V, K166E, I167T, E235K, K162Q, V16I, M153T, M233T, H231P, R168H, E235D, Q204H, H231Q, G232R, I229L | T230A, **Q80R**, Y39F, E235G, D234G, **H231L**, K79R, M233L, T50S, K26R, G232S, Y145C, G232V, A154T, S147C, N146D, F99S, S205A, D190A, P192S, Y237D, H139N, Y151N, K126T, N164I, D133A, L18M, N149K, G4A, N146I, V150A, T108S, V224L, F46L, Y200C, **S65A**, H148N, I167L, V61L, I161T, S65T, **C48S**, Q69L, H148R, V68G, E222G, E222A, Y66H, L207Q | E6A, W57G, Y66F, Y66L, Y66S, G67A, E132D, L141M, S147A, H148D, H148Q, N149C, I167A, H169C, Q183E, Q183R, S202D, T203V, Q204E, S205C, E222Q, A227G, G228A, T230V, M233E, M233G, M233P, D234H, D234S, D234T, E235L, E235Q, L236S | *** |

Shown in ranked order with most enhancing substitutions first according to GMMA. Substitutions in bold are discussed in the main text.
* des11 background mutations not discussed in [19]
** reported to support Trp chromophore [33,34]
*** 84 substitutions omitted

### Appendix 1: IFS detection

Irreversible fatal substitutions (IFS) are ignored during estimation of initial values in steps 1 and 2, because these steps rely on the average effect, $\langle \Delta F \rangle$, which is sensitive to the inclusion of apparently highly destabilizing substitutions. To detect IFS, we estimate the likelihood of the subset of variant data that involves a given substitution using a modified multinomial probability. A substitution is marked as IFS if the distribution of active and inactive *N*-mutants is unlikely, presumably due to many inactive variants, considering that e.g. a double mutant is more likely to be active than an 8-mutant.

The probability parameters of the multinomial function, $\boldsymbol{p} = \{p_{f,N}\}$, of active (f=a) and inactive (f=d) *N*-mutants are estimated using the full data set as $p_{\mathrm{f,N}} = n_{\mathrm{f,N}}/n_{\mathrm{tot}}$, where $n_{\mathrm{tot}}$ is the total number of variants. The variant carrying most substitutions is a 15-mutant which result in 30 probabilities, $p_{a1}$, $p_{d1}$, $p_{a2}$, $p_{d2}$, …, $p_{a15}$, $p_{d15}$. Secondly, a likelihood of the variant set carrying a given substitution is calculated as

$$L(\boldsymbol{x}, n|\boldsymbol{p}) = \mathrm{MN}(\boldsymbol{x}|n, \boldsymbol{p})\mathrm{M}(n|\boldsymbol{p})$$

where $\boldsymbol{x} = \{n_{f,N}\}$ is the distribution of the variants on active and inactive *N*-mutants that sums to *n*, $\mathrm{MN}(\boldsymbol{x}|n, \boldsymbol{p})$ is the multinomial distribution, and the modification $M(n|\boldsymbol{p})$ is the number of possible ***x*** for *n* given ***p*** calculated by the binomial coefficient

$$\mathrm{M}(n|\boldsymbol{p}) = \binom{n+k-1}{k}$$

where *k* is the length of ***p***. This modification is the same as the number of combinations for *k* indistinguishable balls (variants) into *n* distinguishable boxes without exclusion. To keep numbers in a convenient range this is not normalized further. A substitution is identified as IFS if none of the variants that contain the substitution are active (log fluorescence less than 2.7) and $\mathrm{L}(\boldsymbol{x}, n|\boldsymbol{p}) < 0.001$.

The modification, $\mathrm{M}(n|\boldsymbol{p})$, is necessary because we wish to compare multinomial probabilities that are conditioned on different *n*. To illustrate this, consider the multinomial probability of two substitutions; one occurring in *n*=10 variants and one occurring in *n*=1000 variants. The multinomial probability of any realization of 10 active and inactive *N*-mutants, is in general much higher than any realization of 1000 variants simply because there are far more ways to combine 1000 variants. If we wish to compare two such realizations we need to handle the conditioning on *n*, and use the joint likelihood of variant number, *n*, and distribution on active and inactive N-mutants, ***x***.
